# IL-25 induces beige fat to improve metabolic homeostasis via macrophage and innervation

**DOI:** 10.1101/474288

**Authors:** Lingyi Li, Lei Ma, Juan Feng, Baoyong Gong, Jin Li, Tianyu Hu, Zewei Zhao, Weiwei Qi, Ti Zhou, Xia Yang, Guoquan Gao, Zhonghan Yang

**Author notes:** Lingyi Li and Lei Ma have contributed equally to this study. **Correspondent footnote** Correspondence.

## Abstract

Beige fat dissipates energy and functions as a defense against cold and obesity, but the underlying mechanisms remain unclear. We found that the signaling of interleukin (IL)-25 including its cognate receptor, IL-17 receptor B (IL-17RB), increased in adipose tissue upon cold and β3-adrenoceptor agonist stimulation. IL-25 induced the browning effect in white adipose tissue (WAT) by releasing IL-4, 13 and promoting alternative activation of macrophages to regulate innervation, which characterized as tyrosine hydroxylase (TH) up-regulation to produce more catecholamine including norepinephrine. Blockade of IL-4Rα and depletion of macrophages with clodronate-loaded liposomes in vivo significantly impaired the browning of WAT. Obese mice administered with IL-25 were protected from obesity on a high-fat diet and the subsequent metabolic disorders, and the process involved the uncoupling protein 1 (UCP1)-mediated thermogenesis. In conclusion, the activation of IL-25 signaling on beige fat might play a therapeutic potential for obesity and its associated metabolic disorders.

## Introduction

Obesity, which affects approximately 13% adults worldwide, has become a major and pressing global problem. Obesity increases the risk of various metabolic disorders such as hypertension, type 2 diabetes, cardiovascular diseases and cancer [1]. As the high energy-density diet and the sedentary lifestyle cause obesity, interventions have been focused on pathways involving a decrease in energy intake and/or an increase in energy expenditure. Brown adipose tissue (BAT) in neck and interscapular region produces heat and is important for small mammals housed in cold environment. Activation of the uncoupling protein 1 (UCP1)-positive adipocytes would release heat via uncoupled oxidation respiratory chain for ATP synthesis [2]. Therefore, brown adipocytes could be a therapeutic target in the treatment of obesity, although the thermogenic tissue is almost lacking in adults [3]. Besides the classical brown fat, the other thermogenic tissue named beige fat tissue could also be a potential target for treatment of obesity [4]. Although both of the two adipocytes are similar in high expression of UCP1, beige adipocyte differs from classical brown adipocyte in the following characteristics. First, from the perspective of evolution, brown adipocyte derives from a myogenic factor 5 (Myf5) and paired box 7 (Pax7)-positive precursors [5], but beige adipocyte precursors from a Myf5-negative, platelet derived growth factor receptor alpha (Pdgfr-α)-positive cell lineage [6]. Second, beige adipocytes with the browning genes such as *Ucp1* and *Ppargc1a* inducible activated by cold and β-agonist stimulation are mainly located in the subcutaneous and epididymal depot [2, 7, 8], while brown adipocytes are located mainly in neck and interscapular region and express high levels of UCP1. Third, the development of brown adipocytes and beige adipocytes was regulated by different pathways [7, 9]. Hence, brown and beige adipocyte are considered as two distinct cell types, whereas the beige adipose tissue innervation depot shares a coincident shift in the gene expression profile of neurons in stellate ganglion projecting to the brown adipose tissue depot [10].

Adipose tissue is an endocrine and immune organ including different kinds of immune cells such as macrophages, eosinophils, T cells, and B cells [11], and therefore characterizes as a typical circumstance tying organismal metabolism to cellular metabolism and immunity. Crosstalk among adipocytes, recruited immune cells and released cytokines play a key role in maintaining the whole homeostasis. Emerging evidence shows that the immune environment also influences the modulation of adipose tissue, especially in the biogenesis of beige adipocytes. For example, eosinophils and alternatively activated macrophages in adipose tissue are crucial for the cold-exposure-induced browning of WAT [12]. In addition, mast cells may also influence the development of beige fat by sensing cold environment and subsequently release histamine to promote UCP1 expression [13]. Therefore, although underlying mechanisms are not fully understand, the diversity of endocrines and cytokines may play a key role in the modulation of adipose tissue, and the immune cells may stimulate the biogenesis of beige fat [14]. Previously we found that nematode infection stimulated type 2 immunity by releasing macrophage-responsive Th2 interleukins (ILs) including IL-25 [15], and modulated body weight against obesity and associated metabolic dysfunction [16]. To further explore whether the process above is related to thermogenesis or not, we hereby investigated the role of IL-25 in modulating the browning effect of adipose tissue.

IL-25 is a member of IL-17 cytokine family (also called IL-17E) present in various tissues. The receptor of IL-25, IL-17RB, is expressed in various cell types, such as epithelial cells, eosinophils and NKT cells [17]. In addition to its role in mucosal or type 2 immunity, IL-25 has also been involved in the protection high-fat diet-induced hepatic steatosis [18] against the excessive lipid accumulation in the liver, or in regulation lipid metabolism to modulate the body weight via alternatively activate macrophages [19]. Whether IL-25 is involved in the biogenesis of Brown-in-white (Brite)/beige adipocytes and associated metabolic disorders is not known.

Moreover, given that the brown and beige adipose tissue can increase thermogenesis by uncoupling the oxidative phosphorylation with UCP1-upregulation, we also hypothesized that constitutively expressed IL-25 induces beige adipose tissue and improves metabolic homeostasis and thermogenesis, and that enhanced IL-25 production in vivo would protect against insulin resistance. The present study was aimed at investigating 1) the effects of β-agonist or cold exposure on IL-25 expression in the adipose tissue, 2) the effects of IL-25 on Brite/beige adipocytes of the white adipose tissue, 3) the role of macrophage during the process that IL-25 induces the beige fat, and 4) whether IL-25 improves the homeostasis against insulin resistance.

## Results

### IL-25 signaling increased in beige adipose tissue upon β3-adrenoceptor agonist stimulation or cold exposure

To investigate the role of IL-25 signaling in the modulation of beige adipose tissue, we chose physiological and classical Brite models in vivo. First, after administration of β3-adrenoceptor agonist (CL-316, 243 (CL)) to 8-week old C57BL/6J mice, a robust browning phenotype characterized as high expression of UCP-1 and multilocular adipocytes in adipose tissue was found (Figure 1A-1C, S1A-S1C). Time-course analysis showed that CL induced the expression of IL-25 and its cognate receptor, IL-17RB, in subcutaneous WAT (scWAT) (Figure 1A and 1D) and epididymal WAT (eWAT) (Figure S1A and S1D). The expression of IL-25 was further confirmed to be induced by CL in scWAT (Figure 1E, left panel) and eWAT (Figure S1E left panel) by ELISA. Figure 1J (left panel) shows that higher IL-25 in adipose tissue did not increase circulatory IL-25.

**Figure 1.**
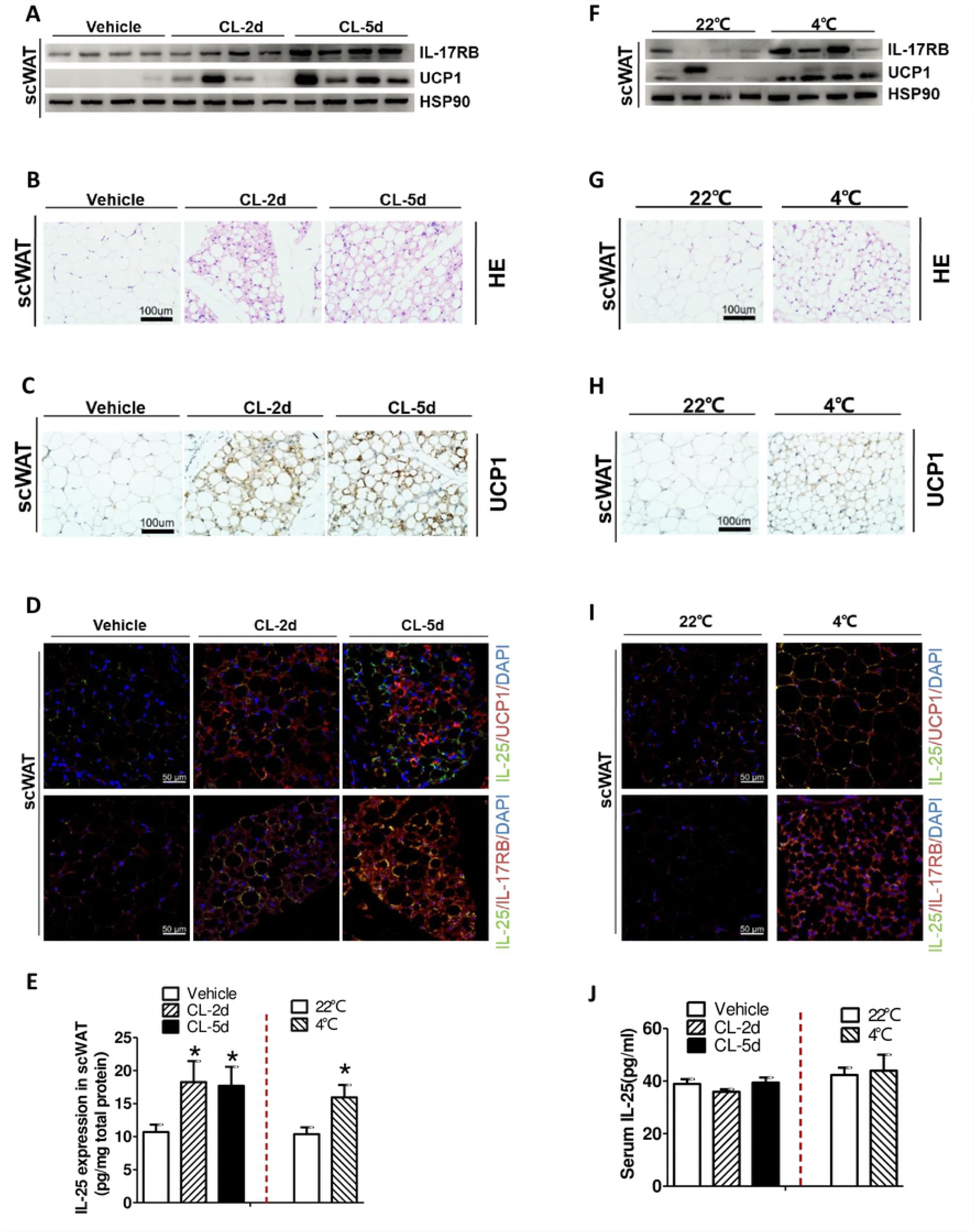
IL-25 signaling increases in subcutaneous beige adipose tissue induced by β3-adrenergic agonist stimulation (A-E) or cold exposure (F-J). Wild type mice were injected with CL (1mg/kg body weight) for 2 (CL-2d) and 5 days (CL-5d), or were placed at controlled temperature (22°C) and cold challenge (4°C) in independently cages for 48h (n=4-5 per treatment). (A and F) The protein level of IL-17RB andUCP1 was analyzed by Western-blotin scWAT. HSP90 was used as a loading control. (B and G) Hematoxylin and eosin (H&E) staining of scWAT (400× magnification). (C and H) Immunohistochemical staining for UCP1of scWAT (400× magnification). (D and I) Immunofluorescent staining for IL-25 (IL-25+ green) and UCP1 (UCP1^+^ red) or IL-17RB (IL-17RB+ red) of scWAT. Nucleus stained with DAPI (blue). Images were photographed at 200× magnification. (E and F) IL-25 protein expression in scWAT (E) or serum (J) (n=4-5 per treatment). *p < 0.05 by two-sided unpaired t-test. Data present as mean ± SEM.

Next, wild type mice were separated into two groups and housed at 22°C or 4°C, respectively, for 2 days. Cold exposure induced a comparable beige phenotype in scWAT characterized as up-regulation of UCP1 (Figure 1F-1H). However, such elevation of UCP1 was not observed in eWAT (Figure S1F-S1H), suggesting that cold-induced sympathetic activation preferentially stimulates the browning in scWAT only. Similarly, the expression of IL-25 and IL-17RB was examined at different temperature. Concomitant to the browning of WAT, cold acclimation also preferentially resulted in an increase of IL-17RB (Figure 1F and1I) and IL-25 (Figure 1E right panel) in scWAT. As expected, cold exposure did not increase circulatory IL-25 (Figure 1J right panel). Unexpectedly, cold exposure only increased IL-25 and IL-17RB in scWAT, did not change in eWAT (Figure S1F, S1I, S1E right panel and S1J left panel), and decreased in the liver (Figure S1J right panel).

### Administration of IL-25 induced the browning of scWAT and repressed DIO chronic low-grade inflammation in vivo

To further investigate whether pharmacologic treatment of IL-25 would induce thermogenic gene expression and browning of WAT, recombinant IL-25 was intraperitoneally administered to normal control mice for 7 days. Figure 2A shows that IL-25 increased the expression of UCP1 protein in scWAT, but not in BAT, in a dose-dependent manner. As the expression of UCP1 also increased after a high dose of IL-25 (1000ng/d) administered in eWAT (Figure 2A), the dose of 1000ng/d for IL-25 was applied in the following experiment. Furthermore, to validate the activity of recombinant IL-25, its receptor, IL-17RB, was also examined and shown to be up-regulated (Figure 2B). Western-blot and RT-qPCR analysis showed that IL-25 induced the expression of UCP1 (Figure 2B and2C). Histologic analysis showed that beige fat was characterized as high expression of the specific mitochondrial UCP1 and multilocular morphology in scWAT and eWAT after IL-25 treatment (Figure 2D and 2E). These data suggested that IL-25 stimulated the development of beige fat.

We next analyzed whether IL-25 could suppress the expression of pro-inflammatory genes and increase the expression of anti-inflammatory genes in diet-induced-obesity (DIO) chronic inflammatory adipose tissue. Figure 2F shows that IL-25 promoted the expression of anti-inflammatory cytokines such as IL-10 and IL-13 and decreased the expression of pro-inflammatory cytokines such as IFN-γ, IL-1β and IL-6.

Next, to further explore the mechanism of IL-25-induced beige, recombinant IL-25 was applied to treat the differentiated 3T3-L1 MBX adipocytes in vitro. As CL might induce the expression of thermogenic or β-oxidation genes in adipocytes in vitro, it was used as a positive control. Figure 2G and 2H show that IL-25 did not augment the expression of thermogenic or β-oxidation genes such as Ucp1, Pgc-1α, Acsl1 and Acox1 at various doses of treatment.

### IL-25 increased the level of IL-4 to stimulate the browning of adipose tissue via inducing alternatively activated macrophages

Figure 3A shows that IL-25 induced the macrophage phenotype switch from pro-inflammatory classically activated macrophages (iNOS as marker) to anti-inflammatory alternatively activated macrophages (ARG1 and YM1 as marker). IL-25 also increased the expression of IL-4 and IL-13 (Figure 3B and3C), which is important for induction of alternative activation program in adipose tissue. However, these changes were not found when treated with irisin (Figure 3D), a myokine that induces the browning of adipose tissue via actions on adipocytes [20].

As IL-25 may induce alternative activation of adipose tissue macrophages and promote IL-4 expression, to explore whether the effects of IL-25 and IL-4 promote the commitment of APs to the beige fat lineage, APs were isolated from scWAT, pretreated with IL-4 or IL-25, and then studied their effects on beige adipocytes. Figure 3E shows that the IL-4-pretreated-Aps, rather than the IL-25-pretreated-Aps, enhanced the expression of UCP1. While APs were co-cultured with peritoneal macrophages treated with IL-25, UCP1 expressed in the APs and then repressed in the presence of IL-4Rα neutralizing antibody (Figure 3F). Next, anti-IL-4Rα antibody was intraperitoneally injected into IL-25-treated normal control mice. Disruption of IL-4/IL-13 signaling with IL-4Rα neutralizing antibody decreased the IL-25-induced mRNA expression and alternatively activated macrophages marker genes in adipose tissues (Figure 3G). Western-blot and RT-qPCR analysis of adipose tissues showed that the expression of UCP1 were increased in WT mice treated with IL-25, whilst blockade of IL-4Rα blunted the expression of UCP1 (Figure 3G and 3H).

### IL-25 regulated adipose tissue innervation by macrophages

As blockade of IL-4Rα by its antibody did not entirely inhibit IL-25-induced expression of UCP1 (Figure 3G and 3H), we explored alternative mechanisms, i.e., enhancing innervation of scWAT and eWAT. Figure 4A and 4B show that expression of tyrosine hydroxylase (TH) was induced after IL-25 administration. Histological staining of TH in scWAT and eWAT showed more sympathetic axons in the IL-25-treated mice than those in the PBS-treated mice (Figure 4C). Similarly, IL-25 also increased the level of norepinephrine in scWAT and eWAT (Figure 4D). These results showed that the IL-25 injection significantly enhanced innervation of scWAT and eWAT.

To further investigate whether brown-adipose-tissue macrophages regulate tissue innervations, we applied clodronate-loaded liposomes to deplete macrophages in DIO mice. Figure 4E shows that depletion of macrophages in DIO mice clearly blunted the IL-25-induced expression of TH and UCP1.

### IL-25 improved the metabolic homeostasis on DIO mice

To investigate whether IL-25 could improve obesity and related metabolic disorder in DIO mice, we firstly fed the C57/BL6J mice with high-fat diet for 12 weeks. After that, the DIO mice were subsequently administered with different doses of IL-25 for another 14 days. At the same time with the administration of IL-25, IL-4 was used as a positive control with a dose of 1μg/mouse, because of its effect on inducing the browning of adipose tissue in HFD-fed mice. Figure 5A shows that IL-25 increased the expression of UCP1 protein in a dose-dependent manner in scWAT and eWAT of DIO mice.

Figure 5B shows that the expression of both UCP1 and TH decreased in DIO adipose tissue, but increased after treated with IL-25, and the change in both UCP1 and TH was only found in scWAT and eWAT, but not in BAT (Figure 5B). The browning effects were shown in the histologic images of scWAT and eWAT, in which adipocytes share biochemical (e.g., high expression of the specific mitochondrial UCP1) and morphological (e.g., high mitochondrial content and multilocular lipid droplet) with BAT in HFD-fed mice treated with IL-25 (Figure 5C and 5D, Figure S2A and S2B). RT-qPCR analysis revealed that administration of IL-25 induced the expression of the thermogenic genes, including *Ucp1*, *Ppargc1α*, *Dio2*, and *Acox1* in HFD-fed mice (Figure S2C and S2D).

Next, to determine whether IL-25 could improve homeostasis against obesity and related metabolic dysfunction via beige fat, total body mass, adipose tissue, GTT and ITT, were examined. Figure S3 shows that beige adipose tissue induced by cold exposure improved the metabolic homeostasis. Those HFD-fed mice treated with IL-25 significantly gained less weight (Figure 5E) and fat mass of eWAT (Figure S2E) compared with those without treatment of IL-25. The adipocyte size of scWAT and eWAT was also decreased (Figure S2F). HFD-fed mice treated with IL-25 also showed lower levels of liver weight and lipids than those without treatment (Figure S2G). Furthermore, a concomitant decrease in blood glucose in the HFD-fed mice was found after IL-25 administration (Figure 5F). Figure 5G and 5H show that, in the HFD-fed mice, those with IL-25 treatment significantly improved glucose disposal and insulin sensitivity compared with those without treatment. As liver, eWAT and muscle from NCD- or HFD-fed mice might reduce p-AKT activity stimulated by insulin, those tissues in mice treated with IL-25 restored the decrease (Figure 5I).

### Macrophage depletion and UCP1 knockout ameliorated IL-25-mediated improvement in glucose homeostasis

To investigate whether IL-25 regulates its anti-obesity and anti-diabetic effects through macrophages, we used DIO mice which were administered with clodronate-loaded liposomes to deplete macrophages in adipose tissue without F4/80, a marker of macrophage, expression (Figure S4A). Figure 6 shows that the stimulatory effect of IL-25 on anti-diabetic was largely depended on macrophages. IL-25 did not reduce fasting blood glucose (Figure 6A), or improve glucose tolerance (Figure 6B) and insulin sensitivity (Figure 6C) in DIO mice treated with clodronate-loaded liposomes. However, IL-25 reduced body weight gain (Figure S4B) and eWAT mass (Figure S4C), lowered liver weight (Figure S4D) in DIO mice as effective as in DIO mice treated with clodronate-loaded liposomes, suggesting that the effect of IL-25 against obesity does not require macrophages. The results above demonstrated that the anti-diabetic effect of IL-25 was dependent on macrophages, while its anti-obesity effect was not.

To investigate whether IL-25 protect from obesity and metabolic associated disorder in DIO mice by stimulating the development of beige fat, wild type (UCP1^+/+^) and UCP1-null (UCP1^−/−^) mice were fed with high-fat diet for 12 weeks and then administered with IL-25 (1μg/mouse) or vehicle for 14 days. During the high fat feeding, no detectable difference between UCP1^+/+^ and UCP1^−/−^ mice in body mass was found (Figure S4E). Genetic ablation of UCP1 may not influence IL-25-mediated anti-obesity effect, because IL-25 reduced body weight gain (Figure S4F) and eWAT mass (Figure S4G), lowered liver weight (Figure S4H) in obese UCP1^+/+^ as effective as in obese UCP1^−/−^ mice. IL-25-mediated improvement in glucose clearance was abrogated in obese UCP1^−/−^ mice, as IL-25 did not reduce fasting blood glucose (Figure 6D), or improve glucose tolerance (Figure 6E) and insulin sensitivity (Figure 6F) in obese UCP1^−/−^ mice. These results indicated that UCP1 played an important role in IL-25-mediated anti-diabetic effect but not in effects on lowering body weight.

## Discussion

In this study, we identified that IL-25 signaling including IL-25 and its receptor IL-17RB increased in mice beige adipose tissue induced by β3-adrenoceptor agonist and cold exposure. Administration of IL-25 promoted the browning of WAT associated with significantly less weight gain and improved glucose and insulin tolerance in HFD-fed obese mice. Importantly, IL-25 may induce beige fat and enhance adipose tissue thermogenesis by alternatively activating macrophages to increase the level of catecholamine, excluding the simple and direct effect of IL-25 on adipocytes. These data provide novel evidence that macrophages were involved in the development of beige fat, and IL-25, serving as a cytokine that connects macrophages and thermogenesis in mice, may play a potential therapeutic role in obesity and associated metabolic disorders.

Our study showed that both β3-agonist (1mg/kg body weight for 2 days) and cold exposure (4°C for 2 days) led to a robust browning phenotype as well as activating IL-25 signal in scWAT, whereas in eWAT only β3-agonist led to the beige effect and cold exposure did not. Jia R. and colleagues had demonstrated the differential responses to cold-induced changes of UCP1 across discrete BAT and WAT depots, which supported the notion that the effects of short-term cold exposure increased thermogenic capacity of BAT, as well as browning of scWAT and, to a lesser extent and later, eWAT [21]. Therefore, the white-to-brown induction of IL-25 firstly appear in the easily beige scWAT such as inguinal WAT. Because cold exposure failure to induce beige in eWAT in a short time (< 2 days) and the difference exist between scWAT and eWAT in the aspect of thermogenesis, the discrepancy of IL-25-induced beige fat also showed the differential responses to IL-25 across discrete BAT and WAT depots. Though IL-25 signaling participated the development of beige fat from white adipose tissue, sharing the biochemical (e.g., mitochondrial biogenesis and high UCP1) and morphological (e.g., robust OXPHOS immunostaining and multilocular lipid droplets) characters with BAT [22], BAT UCP1 level is unchanged, which suggests that IL-25 could not affect the function of BAT. In addition to stimulating thermogenesis, IL-25 can suppress the expression of pro-inflammatory cytokines. Previous studies had demonstrated that the browning of WAT was closely related with changes in the expression of inflammatory genes [23] and obese mice displayed higher expression of pro-inflammatory genes [24].

Our study comprehensively explored the mechanism involving in the process of IL-25 inducing beige fat. IL-25 is considered to be different from other peptide-like inducers, such as irisin that directly acts on adipocytes to stimulate the development of WAT [25]. We directly applied IL-25 on APs, but it could not promote differentiation of APs to beige adipocytes. Using IL-25 to pharmacologically activate alternatively activated macrophages in normal control mice, we were able to place alternatively activated macrophages to the environment of IL-25-induced browning WAT. When APs were co-cultured with peritoneal macrophages, IL-25 could promote APs differentiation towards beige adipocytes via macrophage. Importantly, this phenomenon was disappeared when using IL-4Rα neutralizing antibody in the co-culture medium, suggesting that IL-25 promote APs towards beige depend on IL-4 and IL-4Rα. IL-4Rα is an essential receptor for IL-4/IL-13 inducing alternatively activated macrophages [26], and IL-4 can activate the IL-4Rα signaling in adipocyte precursors (APs) and promote the APs to the beige fat lineage [27]. We validated the fact that IL-25-induced alternatively activated macrophages released IL-4, and then activated the IL-4Rα signaling in APs and promoted the APs to the beige fat lineage in vitro. On the contrary, blockade of IL-4Rα in vivo significantly blunted IL-25-induced expression of UCP1. Collectively, these data suggest that IL-25 regulates the browning of WAT via macrophages and APs involving with IL-4 signal pathway.

We found that blockade of IL-4Rα could not entirely inhibit IL-25-induced expression of UCP1, suggesting that IL-25 also might participate the development of beige fat with other mechanism. The fact that scWAT characterizes as prominent regional variation in beige fat biogenesis with localization of UCP1^+^ beige adipocytes to areas with dense sympathetic neurites, and the density of sympathetic projections dependent on PRDM16 in adipocytes, provides another potential mechanism underlying the metabolic benefits mediated by PRDM16 and UCP1 [28]. The development of beige fat requires noradrenergic stimulation from sympathetic nerve system [29]. Norepinephrine (NE) production by axons that express tyrosine hydroxylase (TH) has been shown to be important for this process [30]. Under the circumstance of cold exposure, sensing cold stimulation by hypothalamus results in enhanced sympathetic nerve branch to induce the browning of WAT [30]. TH mainly present in the cytosol and in some extent in the neuron plasma membrane, catalyzes the rate limiting step in the initial reaction for the biosynthesis of catecholamines such as NE packing in vesicles and exporting through the synaptic membrane.

Norepinephrine released through innervation has been shown to be important for the browning of WAT. This study identified that IL-25 could enhance the sympathetic innervation and then induce the expression of NE in WAT. It was reported that alternatively activated macrophages synthesize NE to sustain adaptive thermogenesis [31]. Whether alternatively activated macrophages are involved in the production of catecholamines to sustain adaptive thermogenesis seem controversial, for that alternatively activated macrophages were reported not to synthesize catecholamines or not to contribute to adipose tissue adaptive thermogenesis [32]. We directly applied IL-25 on peritoneal macrophages to induce alternatively activated macrophages, but failed to induce the expression of TH. This result suggested that IL-25 could not produce NE by alternative activation of macrophages. It was recently illustrated that brown-adipose tissue macrophages controlled tissue innervation [33]. In this study, we administered IL-25 to DIO mice with macrophage depletion by clodronate-loaded liposomes, and found that absence of macrophages impaired IL-25-induced sympathetic nerve branching and browning of WAT. Therefore, we considered that IL-25 regulated WAT innervation and released catecholamines such as NE to induce beige fat by macrophages. Although something is still unknown, modulation of the immune environment in adipose tissue by special cytokines can stimulate the biogenesis of beige fat. For example, accumulation and activation of type 2 innate lymphoid cells (ILC2s) by IL-33 could induce the browning of white adipose tissue [27, 34].

After the effect of IL-25-inducing beige fat had been investigated, the role of which on the thermogenesis and metabolic homeostasis was further explored. In this model, IL-25 decreased body weight gain, improved glucose disposal and insulin sensitivity. The effect of IL-25-mediated improvement in glucose clearance showed a marked dependence on macrophages. IL-25 failed to improve glucose tolerance and insulin sensitivity in DIO mice with macrophages depletion. And the anti-obesity effect of IL-25 did not require macrophages for that depletion of macrophages could not impaired IL-25-mediated lowering of body weight. The data from UCP1^+/+^ and UCP1^−/−^ mice clearly indicated that IL-25 required UCP1 to promote glucose clearance but not in its role of anti-obesity via lipolysis. For the anti-obesity action of IL-25, it was possible to note some lipid metabolism related enzymes, such as lipolytic enzymes (ATGL, MAGL, p-HSL) increased and lipogenic enzymes (ACC) reduced [19]. Though IL-25 could induce the browning effect, but it was noted that lipolysis in brown adipocytes was not essential for cold-induced thermogenesis in mice [35].

In conclusion, our results demonstrated that IL-25 induced beige fat via macrophage, improve the homeostasis, and decreased glucose disposal and insulin resistance. Our study indicated that the activation of IL-25 signaling might play a potential therapeutic role against obesity and its associated metabolic disorders.

## Materials and methods

### Animals and In Vivo Experiment

Wild-type C57/BL6J mice were purchased from the center of laboratory animal of Sun Yat-sen University. UCP1^−/−^ mice with C57/BL6J genetic background were purchased from the model animal research center of Nanjing University. All mice were maintained under 12-hr light-dark cycles with a designed environmental temperature (21°C ± 1°C). Four-week-old male mice fed normal control diet (NCD) or high fat diet (HFD-60% kcal fat diet) for 12 weeks to render mice obesity. Except HFD-fed mice, 8 weeks old-male mice were used in all experiments. For cold exposure experiments, C57/BL6J mice were kept under controlled temperature (22°C) at first and then moved to 4°C in independently cages for 48h. For β3-adrenoceptor agonist treatment, C57/BL6J mice were intraperitoneally (i.p.) administrated with CL-316,243 (1mgkg^−1^, Tocris) or vehicle once daily for indicated time points. For IL-25-inducing browning experiments in NCD- or HFD-fed mice, various dose of IL-25 (Biolegend) or vehicle were injected i.p. once daily for 7 days or 14 days. For irisin-inducing browning experiments in NCD-fed mice, irisin (1μg, PeproTech) or vehicle were injected i.p. once daily for 2 or 5 days. For IL-4Rα neutralization in vivo, 125μg IL-4Rα (Cat # 552288, BD Bioscience) or isotype control Ab (Cat # 554687, BD Bioscience) with or without 1μg IL-25 (Biolegend) were diluted with PBS to volume of 0.3ml and injected i.p. twice a week at day 1 and day 4. To deplete macrophages, clodronate-loaded liposomes or empty liposomes (0.2ml/mouse) were injected i.p. once every two days starting 5 days before administration with IL-25 (1μg/d, Biolegend). After IL-25-inducing browning in HFD-fed mice, glucose (2gkg^−1^, Sigma) or insulin (0.75Ukg^−1^, Sigma) were injected i.p. to perform intraperitoneal glucose or insulin tolerance tests in overnight-fasted or 8 hr-fasted mice. After injection, blood glucose concentration was measured using a OneTouch Ultra Glucometer (Johnson) at designed time points. For insulin signaling experiments, IL-25-injected HFD-fed mice were administrated with insulin (0.5Ukg^−1^) through the inferior vena cava, and then inguinal adipose tissue (iWAT) namely subcutaneous white adipose tissue (scWAT), epididymal white adipose tissue (eWAT), interscapular brown adipose tissue (BAT),liver and muscle were harvested within 10 minutes. Cohorts of ≥ 4 mice per genotype or treatment were assembled for all in vivo studies. All in vivo studies were repeated 2-3 independent times.

### SVF isolation

SVF from scWAT of C57/BL6J female mice at age 4 weeks old. ScWAT were washed with PBS, minced and digested with 0.1% type Π collagenase (Sigma) in DMEM containing 3% BSA and 25μg/ml DNase Ι (Roche) for 30min at 37°C. During the digestion, the mixed solution was shaken with hand every 5 min. The mixed solution was filtered through 70μm cell strainer (Falcon) and then centrifuged at 500g for 5 min at 4°C. The floating adipocytes were removed, and the pellets containing the stromal vascular fraction (SVF) were re-suspended in red blood cell lysis buffer (Sigma) for 5 min at 37°C. Cells were centrifuged at 500g for 10 min at 4°C and the pellets were re-suspended in DMEM medium containing 10% FBS and penicillin/streptomycin (100 units/ml).

### Cell Culture

3T3-L1 MBX cells was purchased from the ATCC. 3T3-L1 MBX cells were cultured and grown to confluence in DMEM supplemented with 10% FBS, penicillin/streptomycin (100 units/mL). Adipocytes (3T3-L1 MBX and SVF) differentiation were induced by the beige adipogenic mixture in 10% FBS DMEM medium containing5μgml^−1^ insulin (Sigma), 0.5mM isobutylmethylxanthine (Sigma), 0.5mM dexamethasone (Sigma), 1nM tri-iodothyronine(T3, Sigma), 125μM indomethacin (Sigma), 1μM rosiglitazone (Sigma). Two days after induction, the media was switch to the maintenance medium containing 10% FBS, 1nM T_3_, 5μgml^−1^ insulin for another 6 days. Various doses of IL-25 (Biolegend) and CL (10μM Tocris) was added when cells reached confluence and sustained through 8 days.

### Macrophage isolation

Macrophages were prompted into the peritoneal cavity by injection of 100% mineral oil (0.5ml, Beyotime). After washing with PBS, macrophages were cultured overnight in DMEM with 10% FBS.

### Real-Time PCR

Total RNA from tissue or cells was extracted with Trizol reagent (Invitrogen). RNA concentration was measured by a NanoDrop spectrometer. 1000ng total RNA was reverse transcribed into cDNA by Prime Script^®^ RT reagent Kit Perfect Real Time kit (TaKaRa). RNA-time PCR analysis using SYBR-Green fluorescent dye (Biorad) was performed with a Biorad CFX 96.

### Histology and Immunohistochemistry

Epididymal, subcutaneous white adipose tissue and interscapular brown adipose tissue were fixed in 4% paraformaldehyde. Tissues were embedded with paraffin and sectioned by microtome. The slides were stained with hematoxylin and eosin (HE) using standard protocol. For UCP1 immunohistochemistry, slides of various tissue were blocked with goat serum for 1h. Subsequently, the slides were incubated with anti-UCP1 (1:1000; ab10983, Abcam) overnight at 4°C followed by detection with the EnVision Detection Systems (K5007, Dako). Hematoxylin (ZSGB-BIO) was used as counterstain.

### Immunofluorescence

For immunofluorescence staining, slides were incubated with rabbit anti-mouse IL-17RB (1:1000; H-40, Santa Cruz), rabbit anti-mouse UCP1(1:1000; ab10983, Abcam) or rat anti-mouse IL-25 (1:1000; MAB1399, R&D) overnight at 4°C, followed by staining with a mixture of secondary antibodies containing an Alex Flour 488-Donkey anti-rat IgG (H+L) (1:200; A21208, Life Technologies) and an Alex Flour 594-Donkey anti-rabbit IgG (1:200; R37119, Life Technologies) for 1h at room temperature. The cell nuclei were counterstained with 4’, 6-diamidino-2-phenylindole (DAPI, Sigma) for 15min at room temperature. The slides were observed with a confocal laser scanning microscope (Carl Zeiss Jena).

### Immunoblot Analysis

Tissues and cells were lysed in RIPA buffer (CST) supplemented with 1mM PMSF (Beyotime). The protein concentration was measured by the KeyGen protein assay kit (KeyGen) and total cellular protein (30ng) were subject to western blot analysis. The protein was transferred to PVDF membrane (Minipore) and incubated with primary antibodies against HSP90 (1:1000; C45G5, CST), UCP1(1:1000 for eWAT, scWAT and cells or 1:10000 for BAT; ab10983, Abcam), TH (1:1000; ab112, Abcam), IL-17RB (1:1000; H-40, Santa Cruz), p-AKT (1:1000; 4060S, CST), AKT (1:1000; 4691S, CST). After incubated with goat anti-rabbit IgG/HRP (1:1000; PI1000, Vector Laboratories) secondary antibody, proteins were detected with chemoluminescence using Immobilon Western HRP Substrate (Merck Millipore) on Imagequant LAS 4000-mini (GE Healthcare).

### ELISA

Catecholamine level was detected using a sensitive ELISA kit (Cat # CSB-E07870m, CUSABIO). Moues IL-25 ELISA kit was purchased from R&D (Cat # DY1399). All measurements were performed with standard manufacture protocol. Adipose tissue (100mg) was rinsed and homogenized in 1ml of PBS. The homogenates were centrifuged for 5 minutes at 10,000 rpm for 10min at 4°C and then the supernate was assayed immediately. All samples were normalized to total tissue protein concentration.

### Statistical Analysis

All data are presented as mean ± SEM. Student’s t-test was used to compare between two groups and one-way ANOVA followed by LSD-t test was applied to compare more than two different groups on GraphPad Prism software. p < 0.05 is considered significant.

## Supplemental information

Supplement information includes Supplemental Experimental Procedures and four figures and can be found with this article.

## Author Contributions

L.L., L.M. designed and performed experiments, interpreted data, and provided intellectual input; J.F. performed macrophage and lipid studies, interpreted data, and provided intellectual input; B.G., J.L., T.H., and Z.W. provided intellectual input; W.Q., T.Z., and X.Y. provided reagents and intellectual input; G.G. direct the study and language revision; Z.Y. conceived and designed the study, performed the experiments and wrote the paper.

## Acknowledgements

This study was supported by National Nature Science Foundation of China: 81370945, 81471033, 81400639, 81570764, 81570871, 81572342, 81600641, 81770808, 81701414; National Key Sci-Tech Special Project of China: 2013ZX09102-053, 2015GKS-355; Program for Doctoral Station in University: 20130171110053; Key Project of Nature Science Foun dation of Guangdong Province, China: 2015A030311043,2016A030311035; Guangdong Natural Science Fund: 2014A020212023, 2014A030313073, 2015A030313103, 2015A030313029; Guandong Science and Technology Project (2014A020212023, 2015B090903063, 2016A020214001); Guangzhou Science and Technology Project: 201508020033, 201510010052, 201707010084, 201807010069, 201803010017; Pearl River Nova Program of Guangzhou Municipality, China, Grant Number: 201610010186; Chang jiang Scholars and Innovative Research Team in University: 985 project PCSIRT 0947; Fundamental Research Funds for the Central Universities of China (Youth Program 16ykpy24, 31610046). The funding agencies had no role in study design data collection and analysis, decision to publish, or preparation of the manuscript.

## Figure legends

**Figure 2. IL-25 induces the browning of WAT and affects inflammatory cytokines in vivo independent of a direct action on adipocytes.** (A) Immunoblotting was used to quantify the expression of UCP1 (3 representative bands are shown) in adipose tissues of WT mice administrated with various doses of IL-25for 7 days (n=5). (B-E) Wild-type mice fed with normal chow (NC) were injected with IL-25 (1μg/day) over 7 days. Immuno-blot analysis of IL-17RB and UCP1 protein in WAT (B). RT-qPCR analysis of UCP1 mRNA in scWAT and eWAT (C). Mice WAT sections from the mice above were stained with H&E (D) and UCP1 (E). 400× magnification. (F) C57/BL6J mice (n=5) fed HFD for 12 weeks were administrated with vehicle or IL-25 (1μg/day) over 14 days and then qPCR analysis of genes associated with pro/anti-inflammatory cytokines in scWAT and eWAT. (G-H) Differentiated 3T3-L1 MBX cells were treated with various doses of IL-25 and CL. Western blotting against UCP1 (G) and mRNA expression of thermogenesis or β-oxidation genes (H).

**Figure 3. IL-25 stimulates IL-4/IL-13 release to promote the browning of adipose tissue via inducing alternatively activated macrophages.** (A-C) WT mice (n=5) were injected with vehicle or IL-25 (1μg/day) over 7 days, and then (A) qPCR analysis of markers associated with macrophage polarization and (B) the mRNA expression of IL-4/IL-13genein WAT. (C)IL-25 stimulates IL-4/IL-13 release in WAT. (D) C57/BL6J mice (n=4-5) were injected with vehicle or irisin (1μg/day) for 3 or 5 days and genes associated with macrophage polarization were analyzed in WAT. (E) UCP1examined by western-blot in differentiated SVF (APs) pretreated with IL-4 (50ng/ml) or IL-25 (50ng/ml). (F) UCP1examined by western-blot in differentiated APs co-cultured with peritoneal macrophages stimulated with IL-25 (50ng/ml) and IL-4Rα neutralizing antibody (50μg/ml). (G-H) C57/BL6J mice (n=5) injected with vehicle or IL-25 (1μg/day) for 7 days and IL-4Rα neutralizing antibody (125μg/day) or isotype control were injected at Day1 and Day4. Real-time PCR analysis of mRNA level of Ucp1, Arg-1, Ym-1 (G) and Western-blot analysis of UCP1 protein level in WAT. (H) HSP90 was used as a loading control. *p < 0.05 by two-sided unpaired t-test. Data present as mean ± SEM.

**Figure 4. IL-25 regulates the innervation of white adipose tissue by macrophages.** (A-D) C57/BL6J mice (n=5) injected with vehicle or IL-25 (1μg/day) for 7 days. (A) Western-blot analysis of TH protein level and (B) real-time PCR analysis of mRNA level of Th in WAT. (C) Representative Immunofluorescence images of TH in WAT. (D) Total norepinephrine production in WAT measured by ELISA. (E) DIO mice (n=5) administrated with clodronate-loaded liposomes to obliterate macrophages and then injected with vehicle or IL-25 (1μg/day) for 14 days. Western-blot analysis of TH and UCP1 protein in WAT. *p < 0.05 by two-sided unpaired t-test. Data present as mean ± SEM.

**Figure 5. IL-25 improves the metabolic homeostasis on DIO mice against obesity and insulin resistance.** (A-I) C57/BL6J mice fed NCD or HFD for 12 weeks (n=5 per treatment) were injected with vehicle and various doses of IL-25 over 14 days. (A) Western-blot analysis level of UCP1 protein in scWAT and eWAT from different treatment mice. (B) Western-blot analysis for level of TH and UCP1 protein in scWAT, eWAT and BAT of HFD mice treated with vehicle or IL-25 (1μg/day) for 14 days. (C-D) Representative images of scWAT (C) and eWAT (D) stained forUCP1. 200× magnification (top), 400× magnification (bottom). (E and F) Changes in body mass (E) and fasting blood glucose (F) in HFD-induced obese mice injected with vehicle or IL-25 for 14 days. (G) Glucose tolerance test (GTT) was conducted by intraperitoneal injection of glucose (2gkg^−1^) and measurement of blood glucose concentration with OneTouch Ultra Glucometer at designed time points in overnight-fasted mice. (H) Insulin tolerance test (ITT) was done by intraperitoneal injected of insulin (0.75Ukg^−1^) and measurement of blood glucose concentration by OneTouch Ultra Glucometer at designed time points in 8 hours-fasted mice. (I) Western-blot analysis of the phosphorylation of AKT and total AKT in eWAT, liver and muscle. The tissues were harvested within 10 minutes after an injection of insulin (0.5Ukg^−1^). #p < 0.05, compared to NCD group, *p < 0.05, compared to HFD group by two-sided unpaired t-test. Data present as mean ± SEM.

**Figure 6. Depletion of macrophages and genetic ablation of UCP1 block IL-25-mediated improvement in glucose clearance.** (A-C) DIO mice (n=5) administrated with clodronate-loaded liposomes to deplete macrophages and then injected with vehicle or IL-25 (1μg/day) for 14 days. (A) Changes in fasting blood glucose. (B) Glucose tolerance test (GTT) was conducted by intraperitoneal injection of glucose (2gkg^−1^) and measurement of blood glucose concentration with a OneTouch Ultra Glucometer at designed time points in overnight-fasted these mice. (C) Insulin tolerance test (ITT) was done by intraperitoneal injected of insulin (0.75Ukg^−1^) and measurement of blood glucose concentration by a OneTouch Ultra Glucometer at designed time points in 8 hr-fasted these mice. *p < 0.05, compared to HFD-control-PBS group by two-sided unpaired t-test. Data present as mean ± SEM. (D-F) Wild type (UCP1^+/+^) and UCP1-null (UCP1^−/−^) mice (n=5 per treatment) were fed with high-fat diet for 12 weeks and then administrated with IL-25 (1μg) or vehicle for 14 days. (D) Changes in fasting blood glucose in these mice. (E) Glucose tolerance test (GTT) was conducted by intraperitoneal injection of glucose (2gkg^−1^) and measurement of blood glucose concentration with a OneTouch Ultra Glucometer at designed time points in overnight-fasted these mice. (F) Insulin tolerance test (ITT) was done by intraperitoneal injected of insulin (0.75Ukg-^1^) and measurement of blood glucose concentration by a OneTouch Ultra Glucometer at designed time points in 8 hr-fasted these mice. *#p <0.05, compared to WT-HFD-IL-25group, *p < 0.05, compared to WT-HFD-PBS group by two-sided unpaired t-test. Data present as mean ± SEM.

